# Focal electrical stimulation on an alcohol disorder model using MRI-compatible chronic neural monopolar carbon fiber electrodes

**DOI:** 10.1101/2022.05.16.492128

**Authors:** Alejandra Lopez-Castro, Diego Angeles-Valdez, Gerardo Rojas-Piloni, Eduardo A. Garza-Villarreal

## Abstract

Neuromodulation interventions, such as Deep Brain Stimulation (DBS) and repeated transcranial magnetic stimulation (rTMS), are proposed as possible new complementary therapies to treat substance use disorders (SUD) such as alcohol use disorder (AUD). It is hypothesized that neuromodulation may induce neural plasticity in the reward and frontostriatal systems via electrical field induction, possibly reducing symptoms. Preclinical self-administration rodent models of AUD may help us gain insight into the effects of neuromodulation therapies on different pathology, as well as the neural mechanisms behind the positive effects. DBS, or any type of brain stimulation using intracranial electrodes in rodents, would benefit from the use of MRI to study the longitudinal effects and mechanisms of stimulation as well as novel targets, as it is a non-invasive technique that allows the analysis of structural and functional changes in the brain. To do this, there is a need for MRI-compatible electrodes that allow for MRI acquisition with minimal distortion of the magnetic field. In this protocol, we present a method for the construction and surgery of chronically implantable monopolar carbon electrodes for use in rats. Unlike conventional electrodes, carbon electrodes are resistant to high temperatures, flexible, and generate fewer artifacts in MRI compared to conventional ones. We validated its use by using a focal electrical stimulation high-frequency (20 Hz) protocol that lasted ~10 sessions. We propose that this technique can also be used for the research of the neurophysiological bases of the neuromodulatory treatment in other preclinical substance use disorders (SUD) models.

## Introduction

Neuromodulation encompasses technologies that apply electrical currents with a variety of parameters, through implanted or non-implanted electrodes, to achieve a functional activation or inhibition of a group of neurons, pathways, or circuits (Krames et al. 2009). The main neuromodulation techniques used in humans are repetitive transcranial magnetic stimulation (rTMS) and deep brain stimulation (DBS) (Klooster et al. 2016). Currently approved by the Food and Drug Administration (FDA), is rTMS therapy for adjunctive treatment for nicotine use disorder, and DBS for essential tremors, dystonia, obsessive-compulsive disorder, and Parkinson’s disease (Edwards et al. 2017). Alas, no consensus exists regarding the mechanisms of action involved in rTMS or DBS (Diana et al. 2017). The effects and mechanisms of neuromodulation in SUDs can be further tested through the use of animal models (Ankeny et al. 2014) which can provide valuable information that could be translated to humans. For instance, using MRI in longitudinal animal models for SUDs treated with neuromodulation, researchers may be able to find neuroimaging biomarkers (Aydin, Unal Aydin, and Arslan 2019), effects, and mechanisms related to clinical outcomes, which could be later translated to human studies (Silva and Bock 2008); (Spanagel 2017). To study focal repeated stimulation in rats, similar to rTMS and DBS, intracranial electrodes are currently a practical solution. However, there are a few major methodological challenges to use intracranial electrodes in longitudinal studies with MRI. Metal electrodes are most commonly used for stimulation in rats, yet are highly susceptible to create artifacts which leads to the loss of signal around the region where the electrode is placed, as well as distortion and lower signal-to-noise ratio (SNR) (Redpath 1998). Carbon fiber electrodes have been used in neuroscience as an option for recording (Joshi-Imre et al. 2019; Chuapoco et al. 2019) and stimulating brain regions (Gallino et al. 2019). These electrodes have proven to be a better choice for the improvement of SNR ratio and MRI artifacts due to their physical properties like high resistance, lightweight, and low density. Physical properties like high resistance, lightweight, and low density. Composed of 900 GPa, with thermal conductivity of 1000 W/mK and electric conductivity of 106 S/m (Zhao et al. 2019). However, the viability of its use in the treatment of SUDs and in longitudinal MRI studies has not been tested.

Therefore, the aim of this work is to provide a simple and low-cost approach for the assembly of chronically implantable monopolar carbon electrodes and their use for repetitive focal stimulation in AUD and we proposed it can be used in other models of SUDs.

## Materials and Equipment

**Table 1.**
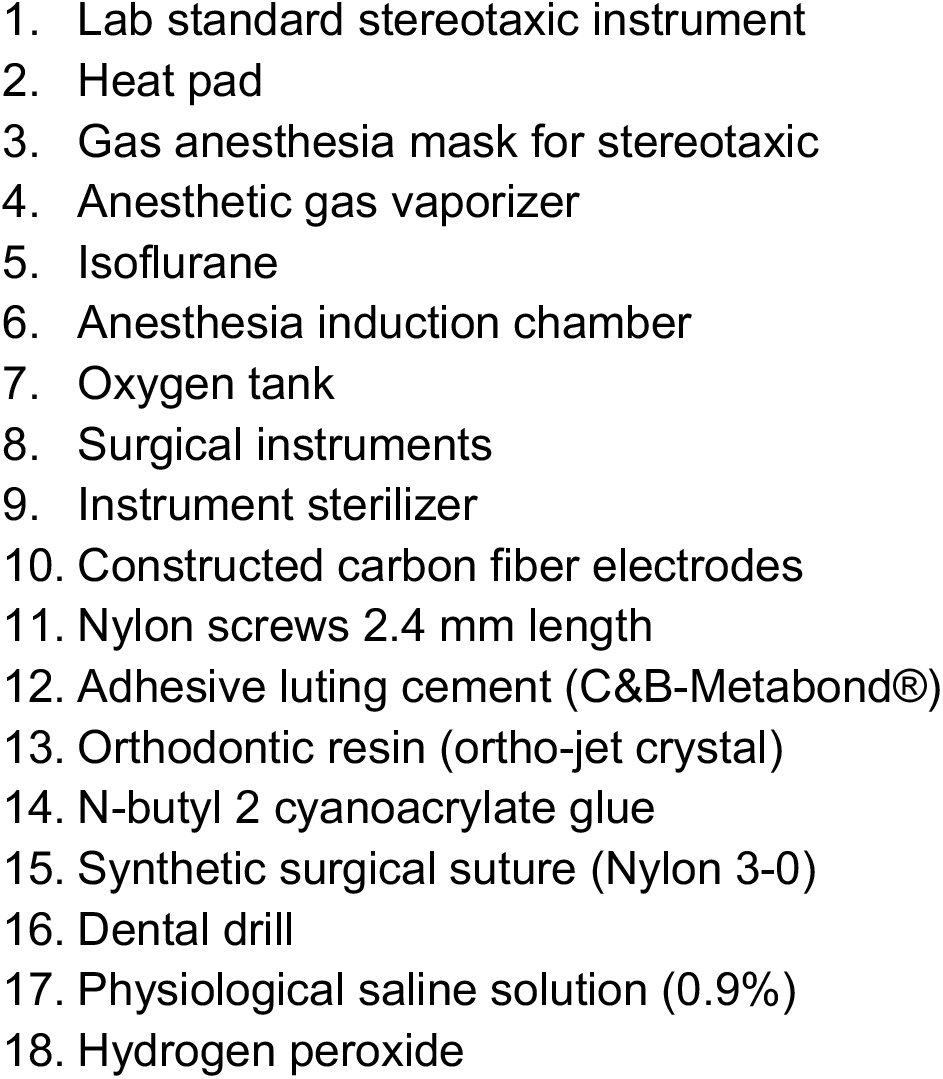
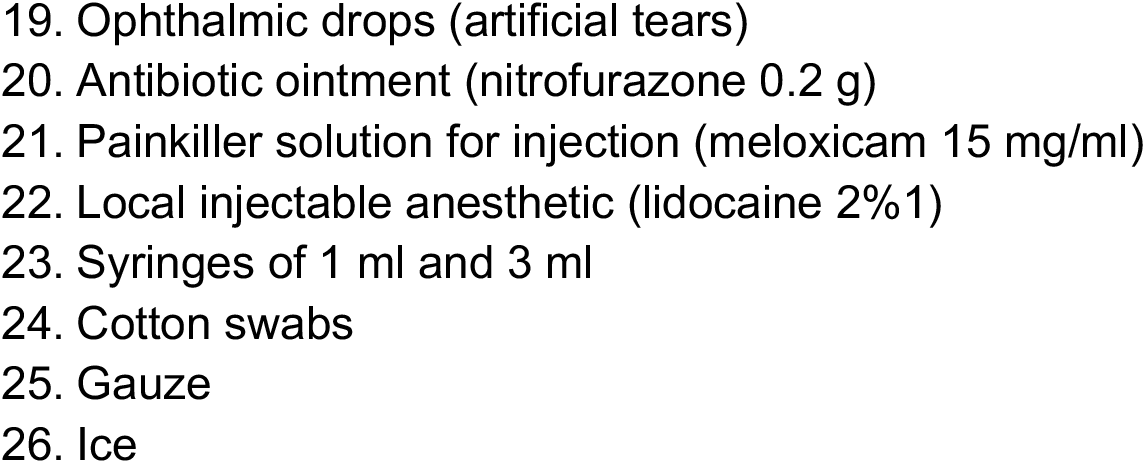
Materials.

**Table 2.**
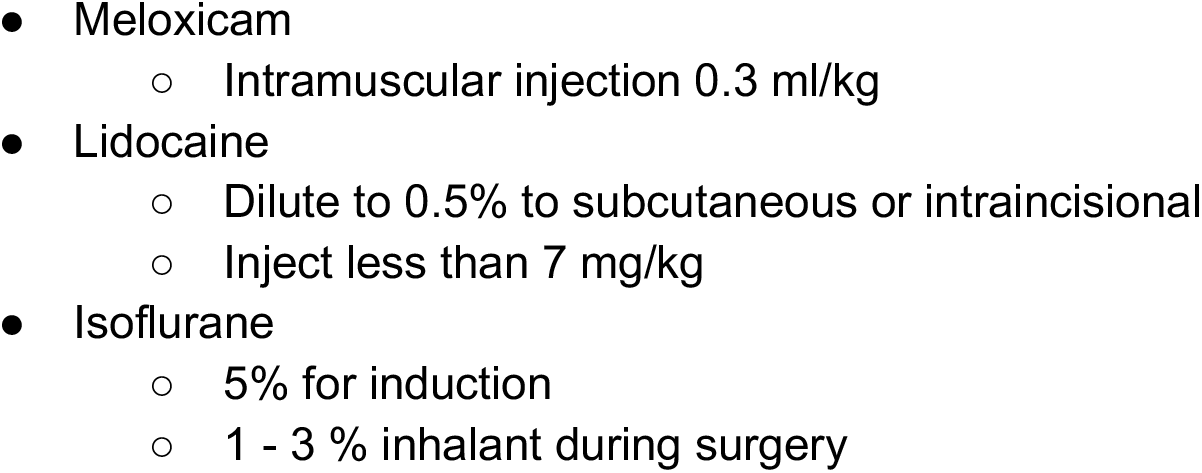
Medication and dosage.

## Methods

The study was approved by the Animal Research Committee of the Instituto de Neurobiología at Universidad Nacional Autónoma de México No. 119-A. All surgical, experimental, and maintenance procedures were carried out in accordance with the Reglamento de la Ley General de Salud en Materia de Investigación para la Salud (Health General Law on Health Research Regulation) of the Mexican Health Ministry that follows the “Guide for the care and use of laboratory animals’’ (National Research Council et al. 2011) and the “Norma Oficial Mexicana” (Mexicana 1999). Also in accordance with the recommendations of the Institute of Laboratory Animal Resources Commission on Life Sciences National Research Council, 1996 and the Directive 2010/63/EU of the European Parliament and of the Council.

### Animals

Twelve adults (male n = 6) and (female n = 6) Wistar rats (*Rattus norvegicus albinus*) were obtained on postnatal day 25 (P25) from the vivarium of the Institute of Neurobiology in Queretaro, Mexico. Animals were individually housed in standard cages in a room with a 12:12 dark cycle/light, controlled temperature (23 °C), and had free access to food.

### Experimental outline

The objective of this work was to validate the use of carbon fiber monopolar electrodes for chronic implantation in an ethanol self-administration model with longitudinal MRI acquisition (Figure 1). Electrode implantation was performed once the *ethanol self-administration model* was established. For this model, we used the *Intermittent access two-bottle choice* (*IA2BC*) (Carnicella, Ron, and Barak 2014; Simms et al. 2008; Wise 1973). Rats were individually housed at P35 and received at least one week of acclimatization with two bottles of water, the same to be used for the IA2BC model and handling. At P45, rats received 24-h sessions of free access to two bottle choices of water and 20% ethanol solution on Monday, Wednesday, and Friday, with 48-h withdrawal periods during the weekends. The placement of the bottles was alternated each drinking session to control for side preferences. During the withdrawal periods, rats received two bottles of water.

**Figure 1.**
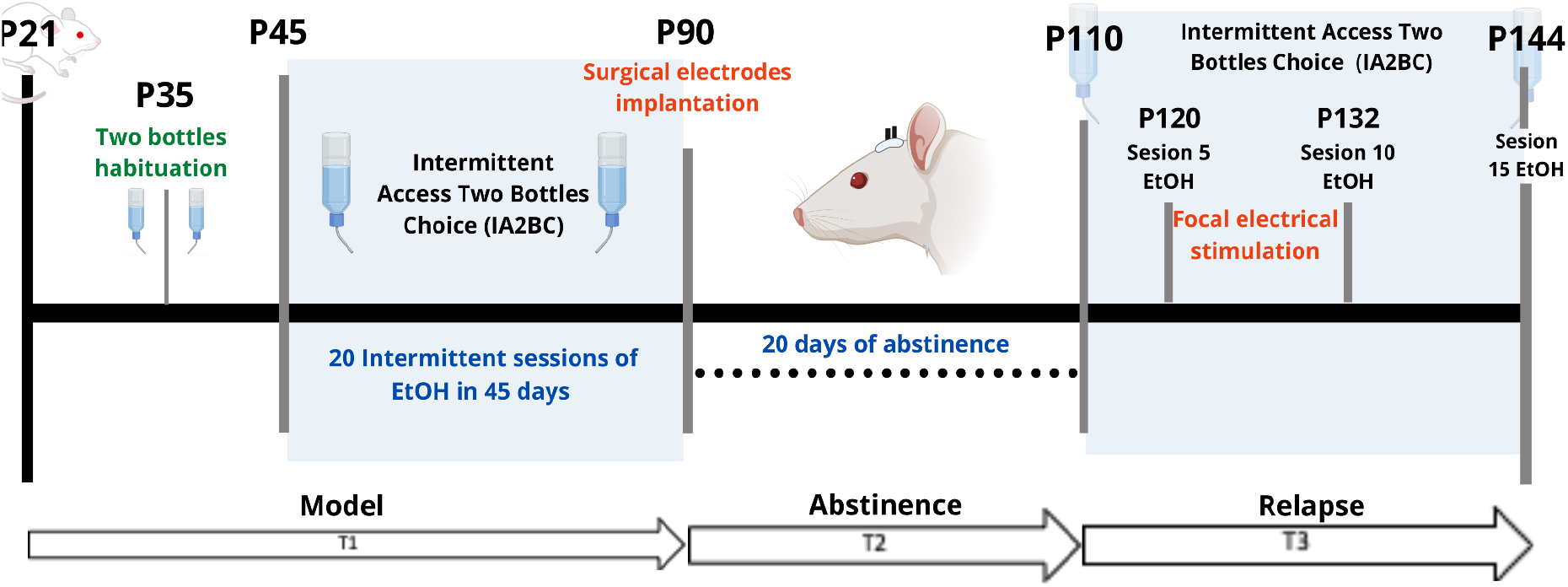
Experimental outline. T1 (Time 1) corresponds to the time encompassing the onset of IA2BC. T2 (Time 2) separates the 20-day period of abstinence. T3 (Time 3) corresponds to the phase of the model where the relapse phenomenon was observed. P = Postnatal age in days. EtOH = 20% ethanol.

As for the timeline, the IA2BC model started on P45 (Time 1 or T1) and lasted for 45 days which included 20 sessions in total, ending in P90. The electrode implantation surgery for stimulation was done at P90, with 10 days of recovery from surgery (Time 2 or T2). Later, at P110, the rats were MRI-scanned (Time 3 or T3). Between P110 and P144 the IA2BC was reestablished with 15 sessions in total to measure relapse and alcohol use. It was during this time that the repeated focal electrical stimulation intervention was applied. T3 was subdivided into 3-time points: pre-stimulation (PreStim), stimulation (Stim), and post-stimulation (PosStim). Before stimulation, we randomly divided the sample into sham (placebo) (n = 6) and active (n = 6) stimulation groups. The sham stimulation group was treated exactly the same way as the active group, except that, during the stimulation sessions, they did not receive any stimulation.

### Carbon electrodes construction

The construction of the monopolar carbon electrodes was based on the Gallino et al., (2019) design. The electrodes were constructed using a cortical fiber of 0.28 mm in diameter (Easy Composites, Stroke on Trent, UK #CFROD-028) and an extracranial fiber of 2 mm in diameter (Good Winds, Mount Vernon, WA USA #CS070048). The cortical fiber was isolated with 3 layers of spray rubber (Plasti Dip®) and both of the fibers were joined with a carbon epoxy (Atom Adhesives, Fort Lauderdale, FL, USA, #AA-CARB61), which conducts electricity between fibers. Finally, the resistance of the electrodes was measured with a voltmeter and marked with a range of 2kΩ - 8kΩ.

### Surgical carbon electrodes implantation

At P80 rats were handled once a day for 5 minutes by caressing the top of their heads, where the electrodes will be placed later. This was to acclimate them to electrode manipulation and human-rat interactions. At P90 surgical procedure was performed; the technique was adapted from (Santana-Chávez et al. 2020; Rigalli and Di Loreto 2016; Matsumiya et al. 2012) (Figure 2).

**Figure 2.**
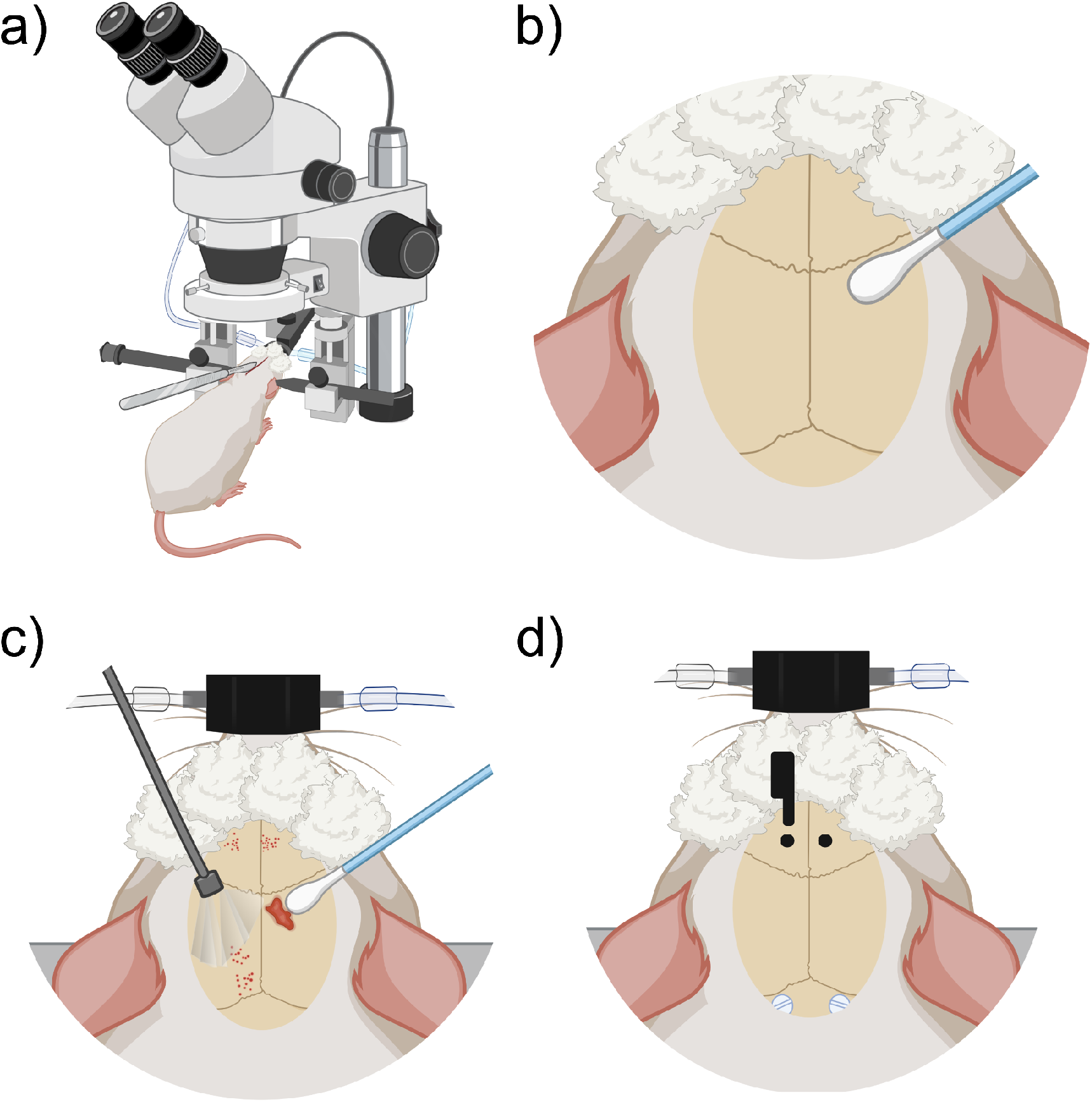
Anesthesia and stereotaxic surgery. a) Rat on base for the stereotactic frame. Rat under anesthesia vaporized with 3% isoflurane, on stereotaxic frame base with head fixation and a heated cushion under the body. b) Preparation of the skull and control of local bleeding. Scalp incision on the midline following the direction of the mid-sagittal suture. c) Curettage of skull aponeurosis and cleaning with swabs. d) Coordinate marking of implantation coordinates of nylon screws and carbon electrodes under the microscope, directed by stereotaxic frame tower. Trepans on the skull are the entry site for the cortical implantation of the electrodes. Also visible are the nylon screws that are placed to keep the fibers fixed to the skull and maintain their chronicity.

#### Preparation for anesthesia (~5 to ~10 min)

Rats were anesthetized with vaporized isoflurane (~5% induction) 50/50 isoflurane/oxygen mixture, administered in an induction chamber. A deep anesthesia state was verified with the absence of withdrawal reflexes to pain. The induction with isoflurane took between 5 to 10 minutes and was maintained during the positioning of the rat in the stereotaxic apparatus. The heart rate was maintained at 60 beats per minute and the respiratory thoracic movement must be watched frequently and checked to be around 20 per minute, and be regular and harmonic on both sides of the thorax. Next, diluted lidocaine to 0.5% was applied subcutaneously at <7 mg/kg, at the zone where the first incision would be made. The rats were injected with intramuscular meloxicam at 0.3 ml/kg to reduce intraoperative inflammation. A heating pad placed on the stereotaxic apparatus base helped maintain the temperature at 25 °C during the surgery. The rat was placed in a supine position maintaining a permeable airway with the aid of the incisor immobilizer bar and the intra-aural position pencils. Artificial tears were then administered to the eyes and then covered with clean gauze.

#### Stereotaxic procedure (~1 hr 30 min)

A midline scalp incision of approximately 2 cm was made using a #20 scalpel blade (Figure 2a). The incision starts posterior to the line of the eyes. Bregma and lambda bony landmarks were exposed. Then the skin was moved to the sides with a self-retaining retractor. A peristome was used to separate the periosteum from the cranium bone. Next, hydrogen peroxide was used to achieve hemostasis with the help of cotton swabs. The importance of this step was to make sure that there was no bleeding and to dry the bone as much as possible (Figure 2b). With the scalpel blade, superficial cuts on the cranium bone were drawn and later washed with saline solution and hydrogen peroxide. Once the cranium was dry and there was no apparent bleeding, one drop of N-butyl 2 cyanoacrylate glue was applied to the exposed surface (Figure 2c).

#### Electrodes placement

Using the tip of the tower of the stereotaxic frame, electrodes were placed with Micropore™ breathable paper tape. Lambda and bregma bony landmarks were measured with the help of the tip of the electrodes and made the adjustment of ≤0.1 mm of a difference between both bony marks. Parallel to lambda, about half a centimeter from the midline, 2 mm diameter circles were marked to place plastic screws (P1tec, Roanoke VA, USA #0-80X3/32N). Subsequently, trepans were made to place the screws. Bleeding control was done with sterile 0.9% saline and cotton-tip applicators (Figure 2d). Finally, the position of the screws was sealed with adhesive luting cement (C&B-Metabond ®).

#### Stereotaxic Coordinates

Under the sight of a surgical microscope, a dental drill was used to make one-millimeter diameter holes bilaterally into the skull at the prelimbic cortex (PrL) (Paxinos and Watson 2006) (Bregma 3.2 AP, 0.4 ML, 3.7 DV) or “area 32” (Paxinos and Watson 2013). The electrodes were placed carefully and slowly, acquiring the DV coordinate 1 μm at a time. Once in place, they were sealed in position with adhesive luting cement, and a layer of resin was applied around and between all the arrays and screws to create a strong head cap fixed to the skull. Once the edges are smoothed and dried, simple stitches with nylon 3-0 closed the wound and left exposed the electrodes’ extracranial fiber.

#### Postoperative care

Before the rat was removed from the stereotaxic apparatus, the vaporized anesthesia was stopped, but the oxygen supply was kept until spontaneous movement appeared. Meanwhile, to manage pain relief, and inflammation, meloxicam (0.3 ml/kg) was intramuscularly injected. A nitrofurazone ointment was applied along the wound, and 1ml of saline 0.9% solution was injected between shoulder blades to maintain the electrolyte balance. The eyes were cleaned and artificial tears were applied once again. When the rat awoke, it was returned to a clean cage, and the bottom of the cage was covered with paper towels to prevent choking or ingestion of the bedding. Each rat was housed individually in cages and monitored closely for 7-10 days after the surgery. If active bleeding was detected, reddened skin, or any signs of discomfort, the nitrofurazone ointment was applied up to 3 times a day. If not, one application per day for 3 days was enough to facilitate healthy scar formation.

#### Stimulation

Between P120 and P132, we began the stimulation protocol (Figure 3) during 10 sessions of 10 min for 10 consecutive days with 100 pulses (duration = 0.2 ms, intensity = 400 μA) at 20 Hz in 10 pulses per train of 2 s, and an inter-train interval of 20 s (Figure 4) (Levy et al. 2007; Gersner et al. 2010). Stimulation was applied by means of GRASS S48 Square Pulse Stimulation connected to a GRASS stimulus isolation unit (SIU) and using metal alligators clips insulated exteriorly with Plasti Dip® and welded to flexible electronic cable 20 AWG. Each rat remained in its individual housing and was habituated for 10 days, prior to treatment, to the connection with the alligator clips.

**Figure 3.**
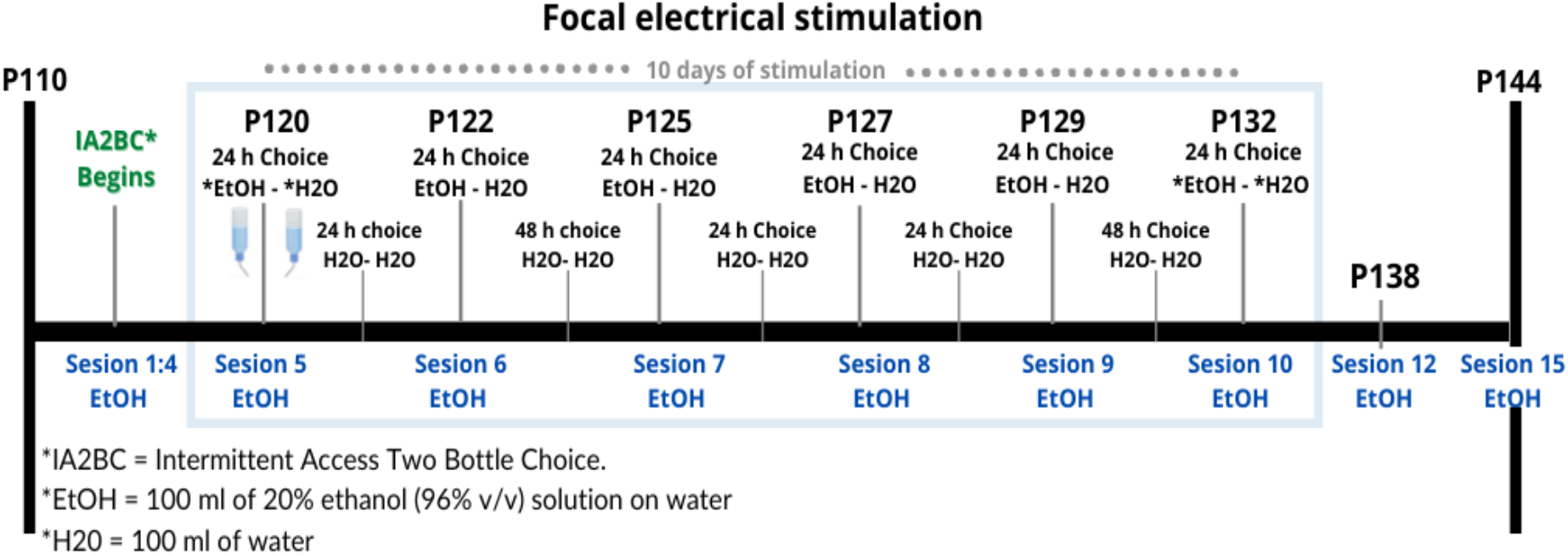
Timing of electrical stimulation and IA2BC during Time 3. Design of the schedule for repetitive focal electrical stimulation consisting of 10 days with a daily session of 10 minutes, this period is shown within the blue rectangle between days 120 and 132 of the age of the rat. In total there were 15 sessions where there was a bottle of water and a bottle of alcohol (EtOH-H2O) to choose to drink, shown in blue. Between each of these sessions, they were offered two bottles of water (H2O-H2O).

**Figure 4.**
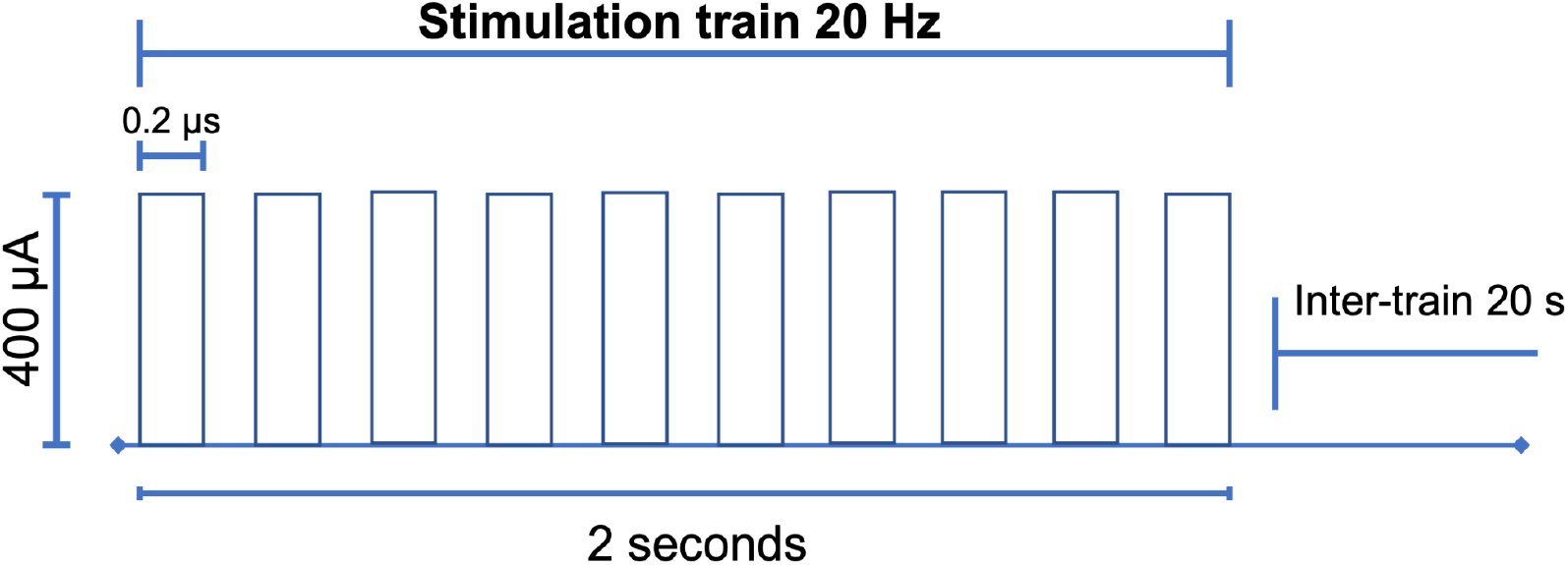
Electrical stimulation design. With the parameters of 100 pulses (duration = 0.2 ms, intensity = 400 μA) high frequency of 20 Hz, in trains of 10 pulses every 2 seconds with an inter-train interval of 20 s. After the choice of the carbon electrode, the research protocol was carried out for 10 consecutive days for 10 min (Levy et al., 2007; Gersner et al., 2010; Moshe et al., 2016).

### Magnetic Resonance Imaging

Magnetic Resonance Imaging (MRI) scanning was done at the Resonance Unit for Rodents and other Animals (URRA), Laboratorio Nacional de Imagenología por Resonancia Magnética (LANIREM) at the Instituto de Neurobiología, UNAM campus Juriquilla, located in Querétaro, Qro, Mexico. Before the acquisition, rats were injected subcutaneously with medetomidine 0.012 mg/kg (Sirmpilatze, Baudewig, and Boretius 2019). During imaging, rats were anesthetized with vaporized isoflurane 5% at induction and 0.5% in a 50/50 mixture of oxygen and vaporized anesthesia. Image acquisition was conducted using a 7T Bruker Pharmascan (Bruker Pharmasan 70/16, US) with a 2×2 surface coil and acquired using Paravision 6.0.1. A 3D Flash sequence T1w with 2 repetitions TR = 30.76ms TE = 5ms and FOV = 25.6 x 19.098 x 25.6 mm and an isometric voxel of 160 microns, was performed at P110 to verify the location of the electrodes and at P144, at the end of the stimulation protocol. All images were converted from Bruker format to nifti using the brkraw tool v0.3.3 (Lee, Ban, and Shih 2020). Anatomical images were preprocessed using an in-house pipeline based on MINC-toolkit-v2 and ANTs which performed the following steps: intensity normalization, center image, denoising, and registering in LSQ6 alignment (https://github.com/psilantrolab/Documentation/wiki/Preprocessing-Rat-Structural-in-vivo). The preprocessing in this manuscript was done for better visualization.

## Results

Twelve rats conserved both stimulation electrodes and remained available for the stimulation protocol for the longitudinal follow-up. To ensure the feasibility of the electrode, we measured electrode resistance between sessions, and all the electrodes held their original values (2kΩ-8kΩ). Additionally, MRI, as a non-invasive technique, was chosen for longitudinal monitoring (Figure 5), and the resulting images were anatomically compared to the Paxinos atlas (Paxinos and Watson 2006).

**Figure 5.**
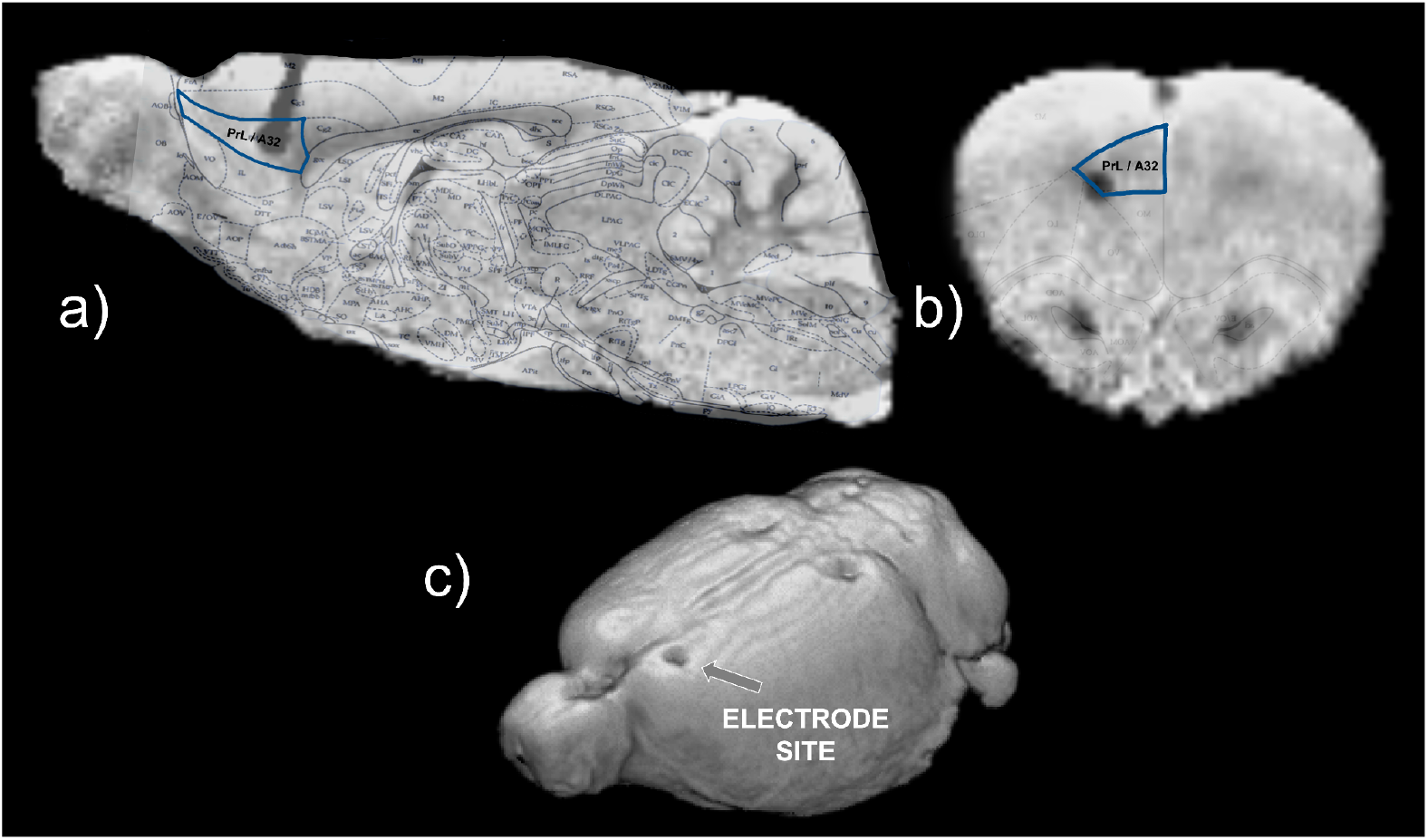
Structural MRI with a reference atlas (Paxinos and Watson 2006) of Rat 12. a) Sagittal slide in blue of the PrL/A32 (prelimbic) region from the atlas and the shadow of the electrode b) Coronal slide in blue PrL/A32. c) 3D model with surface anatomical evidence marks of the electrode implantation in the frontal lobe. The contralateral electrode (right) is not marked since its location is on the surface of the cortex.

Figure 6 shows the results on ethanol intake at all stages of the experiment (T1, T2, T3). The beginning of T3 is also the relapse phase, and all the rats increased their intake in comparison with the baseline. After the stimulation, there were no group differences in alcohol consumption. However, the individual plots showed high variability (Supplementary Table 1) in the individual consumption. Two rats in the sham group, as well as 2 in the active group, reduced their ethanol consumption, while 2 rats in the active group and 1 in the sham group maintained their consumption stable. Two rats in the active group and 3 rats in the sham group increased their ethanol consumption after stimulation. Overall, our results show that 66% of the rats in the active group had a positive effect of stimulation/surgery, while 50% of the rats in the sham group also showed a reduction in consumption.

**Figure 6.**
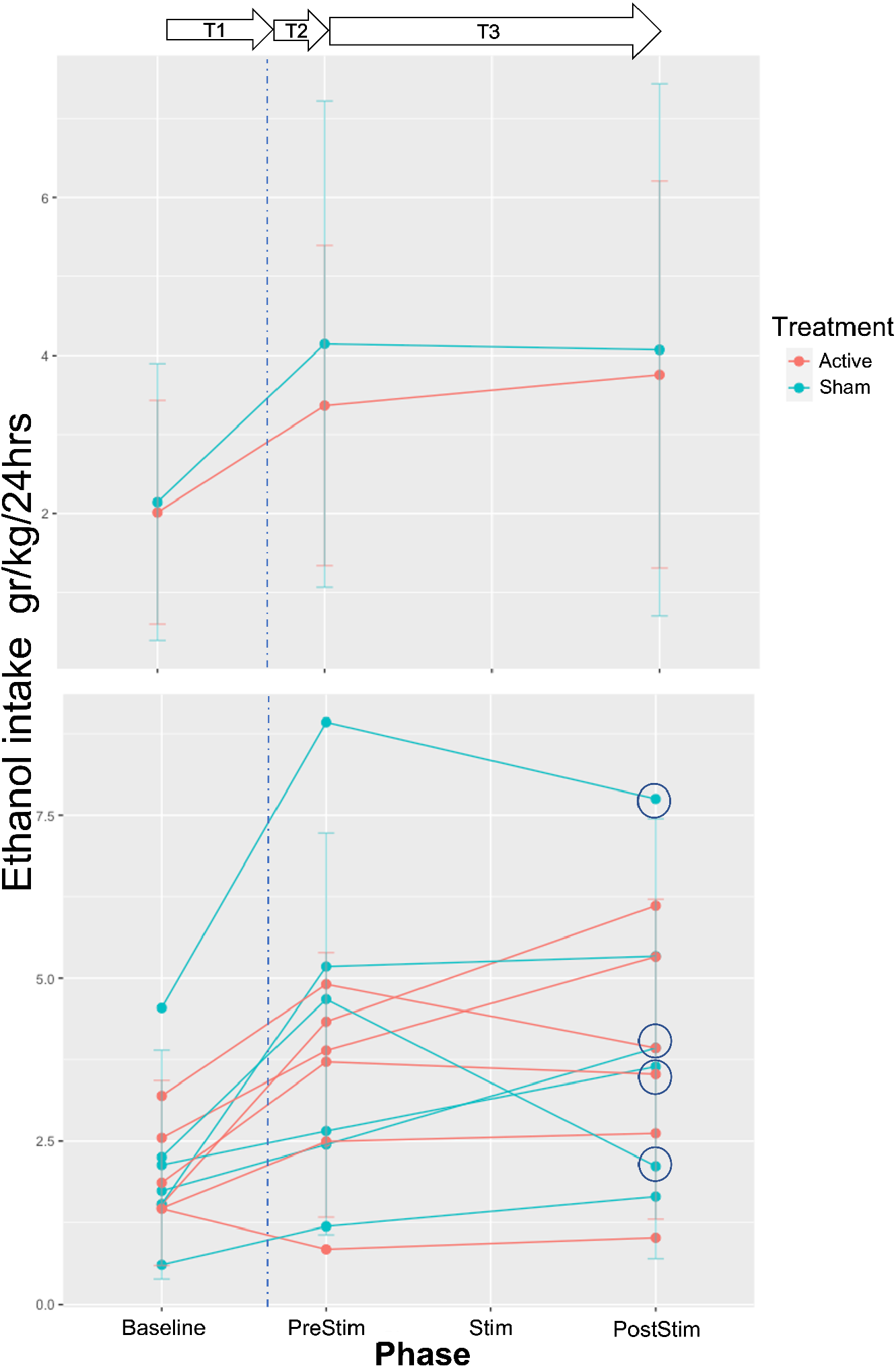
EtOH intake across phases of follow-up. T1 or IA2BC development (baseline) is the 45 days where alcohol is offered in an elective model. The time for the surgery is marked with a blue dotted line (T2 or abstinence). T3 (relapse) comprehends a period before the stimulation treatment (PreStim) and the application of treatment under two conditions: sham/active (Stim) and a period of follow-up after the treatment (PostStim). Circled are the decreasing intakes between PreStim and PostStim evaluation of four rats.

One of our main objectives was that our electrodes would cause only a small lesion in the stimulation area, and with minimal inflammation. Figure 7 shows the longitudinal follow-up of a rat, where there were no structural changes measured with the T1w sequence.

**Figure 7.**
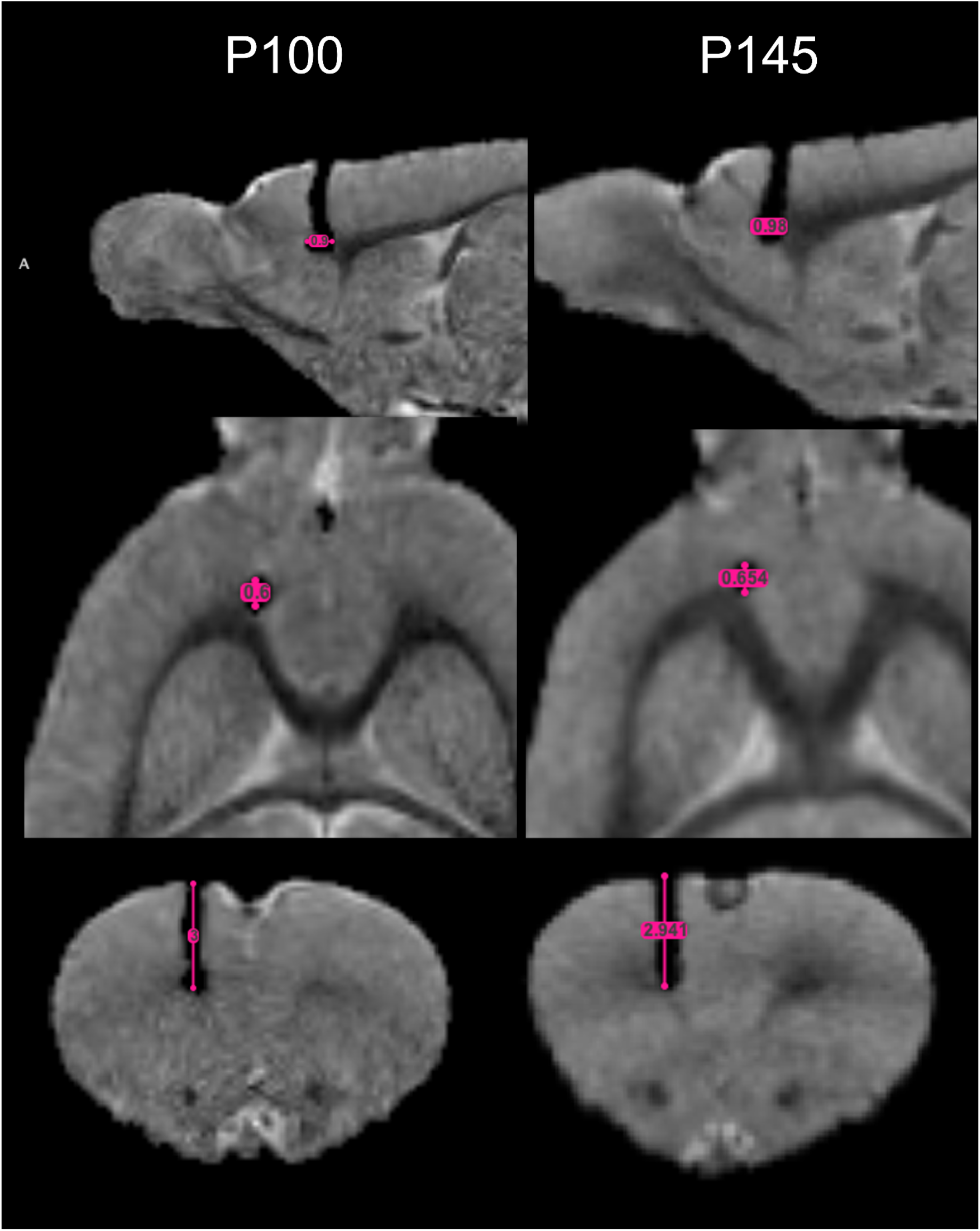
Longitudinal follow-up of rat 12. The implantation surgery was performed at P95 and MRI-scanned at P100. The pink ruler shows the manual measurements in mm of the lesion by the electrode. At P145 the experiment was concluded and an InVivo scan was made for the last time.

## Discussion

Without a doubt it was challenging to assemble electrodes that met the requirements (Geddes and Roeder 2003), of being: 1) MRI compatible, 2) able to perform focal electrical stimulation, 3) well accepted by the body chronically, and also, 4) able to register electrophysiological signals for future studies. Here It described the methods for the construction of carbon monopolar electrodes (Gallino et al. 2019) and proposed their use for the focal electrical stimulation, as an intervention in a preclinical AUD model (McBride and Li 1998). The importance of the intended intervention was to stimulate a brain region that is thought to be the human homolog of the dorsolateral prefrontal cortex, which is the most commonly stimulated target region in rTMS. This region, the prelimbic cortex (PrL) or area 32, is an essential hub of the mesocorticolimbic network (Koob 2013). The nucleus accumbens (NAc) and prefrontal cortex (PFC) PrL, or medial PFC in rats (Laubach et al. 2018) are brian regions that promise to be therapeutic targets for SUDs. Previously, implanted stimulation methods in PFC have decreased alcohol consumption in humans (Voges et al. 2013) and cocaine in rats (Levy et al. 2007). In this work, we stimulated rats that were previously exposed to IA2BC, and we found that four subjects with the greatest consumption of ethanol, decreased their intake in comparison to their consumption at the beginning of relapse or T3. To explain why both sham and active treatments on those four rats had an effect, we agree with Chakravarty et al. (Chakravarty et al. 2016) findings, who proposed that there were changes at the level of synaptic connections in both active and sham stimulation. In their study, active stimulation revealed a remodeling of the structure of the vasculature so that the diameter of the vessels increased. They also found increased volume in the subjects who received stimulation in remote regions, suggesting that neuroanatomical rearrangement also occurs at remote regions connected across multiple synapses (Stone et al. 2011). For future studies, we will analyze the functional and structural effects of stimulation in this AUD model (Bashir, Edwards, and Pascual-Leone 2011), and apply it to other SUD models.

Finally, there were limitations in the model and the stimulation protocol. The sample size is, of course, a problem, however, acquiring a larger sample of rats with higher intake is difficult, as previously described by Carnicella et al., (2014), since only 30-40 % of the rats in the IA2BC model become high drinkers. Nonetheless, we are working on running new experiments to increase the sample size.

In summary, this work describes chronic, MRI-compatible, carbon electrode implantation, and the use of focal electrical stimulation on a preclinical model of AUD with a longitudinal follow-up. Our findings suggest the possibility of decreasing ethanol intake after the stimulation protocol. Further work is needed to elucidate the effects of stimulation in AUD and other SUDs.

## Acknowledgments

Alejandra Lopez-Castro is a doctoral student from the Programa de Doctorado en Ciencias Biomédicas, Universidad Nacional Autónoma de México (UNAM) and has received CONACyT fellowship 1003251. This study was supported by Programa de Apoyo a Proyectos de Investigación e Innovación Tecnológica, Dirección General de Asuntos del Personal Académico (PAPIIT) grants IA202120 and PAPIIT IA201622. The authors would like to acknowledge the vivarium of Instituto de Neurobiología (Dr. Maria Antonieta Carbajo Mata, Dr. Alejandra Castilla León, and MVZ Martin García Servín), and the National Laboratory for Magnetic Resonance Imaging, particularly Dr. Juan Ortiz-Retana for technical assistance. We greatly acknowledge the technical support of Dr. Rafael Olivares. We thank M.S. Alfonso Fajardo-Valdez for his assistance in the MRI visualization. Also, we thank BSc. Diego Ortuzar for the grammatical revision of the manuscript, and Dr. Pavel E. Rueda-Orozco for his recommendations regarding the surgery. Figures 2 and elements from Figure 1 were created with BioRender.com. Finally, we would like to thank Daniel Gallino for his feedback and support with the electrode construction.

## Supplementary Material

**Supplementary Table 1.** Ethanol intake gr/ kg / 24 hrs of the twelve rats.

| Rat | Ethanol intake gr/kg/24 hrs |  |  |
| --- | --- | --- | --- |
| | Mean ± Standard deviation | Max (Min) | Range |
| 1 | 2.42 ± 2.34 | 11.51 (0.28) | 11.23 |
| 2 | 2.02 ± 1.47 | 6.85 (0.18) | 6.67 |
| 12 | 3.27 ± 2.64 | 10.08 (0.26) | 9.82 |
| 16 | 3.42 ± 3.11 | 13.23 (0.00) | 13.23 |
| 17 | 2.56 ± 1.66 | 6.38 (0.39) | 5.99 |
| 18 | 2.48 ± 1.28 | 6.20 (0.39) | 5.81 |
| 25 | 3.67 ± 1.84 | 8.16 (1.22) | 6.94 |
| 26 | 6.07 ± 2.63 | 11.30 (1.67) | 9.62 |
| 33 | 1.16 ± 1.14 | 5.26 (0.12) | 5.15 |
| 39 | 0.87 ± 0.81 | 3.37 (0.14) | 3.23 |
| 45 | 3.25 ± 1.80 | 9.21 (0.21) | 9.00 |
| 48 | 2.83 ± 1.68 | 7.66 (0.63) | 7.03 |

